# Soundscape reflects breeding phenology in colonial seabirds

**DOI:** 10.64898/2026.05.27.728265

**Authors:** Valerie M. Eddington, Danielle T. Fradet, Elizabeth C. Craig, Megan A. Cimino, Easton R. White, Laura N. Kloepper

## Abstract

Migratory seabirds are valuable indicators of marine ecosystem change but can be difficult to monitor during the breeding season due to dense colonies, remote breeding sites, and sensitivity to investigator disturbance. Passive acoustic monitoring offers a minimally invasive alternative to traditional surveys; however, high call overlap in large colonies complicates approaches that rely on identifying individual vocalizations. In this study, we evaluate acoustic energy as a simple soundscape metric for monitoring breeding phenology in colonial seabirds. Using a comparative approach, we deployed autonomous recorders at breeding colonies of Adélie penguins (*Pygoscelis adeliae*) in the Western Antarctic Peninsula and common terns (*Sterna hirundo*) in the Gulf of Maine. We examined seasonal patterns in acoustic energy and compared these trends with known breeding stages and colony observations. Across both species, acoustic energy exhibited distinct seasonal patterns that correspond to key phenological stages, including courtship, incubation, chick rearing, and fledging. These stages are associated with distinctive colony-wide behavioral shifts in colony attendance, territorial interactions, and parent-offspring communication that structure the breeding-season soundscape. Our results demonstrate that colony-wide acoustic energy can capture key phenological transitions in seabird colonies and provide a scalable, minimally invasive approach for monitoring breeding dynamics in remote or rapidly changing environments.

**Highlights:**

- Passive acoustic monitoring can track bioindicator phenology under climate change
- Amplitude captures colony-level activity in dense seabird colonies
- Soundscape patterns correspond to key breeding stages
- Effective in both temperate and polar seabird systems
- Enables scalable, low-disturbance monitoring in remote systems

## 1. Introduction

Seabirds are apex predators within pelagic systems and serve as valuable bioindicators of marine ecosystem health (Diamond and Devlin, 2003; Scopel et al., 2019) and climate change impacts (Cimino et al., 2023; Smith and Craig, 2023). Seabirds are the most threatened avian group globally, with roughly half of all species’ populations suspected or known to be declining (Croxall et al., 2012), which emphasizes the need for effective long-term population monitoring. Despite the critical need for these programs, implementing them in seabird colonies presents logistical and methodological challenges. Seabirds are inherently difficult to monitor due to their wide-ranging at-sea distributions (Lascelles et al., 2012) and remote, rugged breeding colonies, which are often difficult to access, making frequent surveys difficult (Field et al., 2005). Traditional visual methods (e.g., point-counts, photographic estimation, and drone surveys) are labor-intensive, subject to observer bias, and spatially and temporally limited (Edney and Wood, 2021; Pérez□Granados and Traba, 2021). Repeated human presence during breeding can also introduce substantial disturbance to colonies (Brosseau et al., 2024), reducing reproductive success and biasing survey results (Carey, 2009; Carney and Sydeman, 1999). As a result, many colonies remain under- or unmonitored (Santos et al., 2018), and existing data often lack the temporal resolution needed to capture critical life-history events (Hutchinson, 1980). Consequently, monitoring approaches capable of collecting accurate, high-resolution data with minimal disturbance to colonies are needed.

One life-history trait of interest to many monitoring programs is phenology, or the timing of biological events, which is an important indicator of ecological responses to climate change. Shifts in migration, breeding, and molt timing have been widely documented across migratory bird species as global temperatures rise (Charmantier and Gienapp, 2014; Gordo, 2007). For migratory seabirds, successful reproduction requires synchronization with seasonal peaks in prey availability, which are increasingly disrupted by warming oceans and sea-ice loss (Buehler and Piersma, 2008; Frederiksen et al., 2004). While some seabird populations have advanced their breeding, others show delays or no change, reflecting spatial variability in climate impacts (Both and Visser, 2001; Frederiksen et al., 2004). Understanding how climate change influences phenology across species and regions is essential for anticipating population-level responses.

While passive monitoring techniques have been employed to collect seabird phenology data, their application remains limited by practical and analytical constraints. Automated time-lapse camera systems have successfully captured nest-level phenological data in the Antarctic Peninsula (Black et al., 2018), producing results comparable to direct observations (Hinke et al., 2018). However, these systems require intensive image classification and consistent nest tracking, and chicks are often obscured during brooding (Hinke et al., 2018). This highlights the need for a more practical and efficient passive approach for monitoring colonial seabirds.

On the other hand, passive acoustic monitoring (PAM) using autonomous recording units has revolutionized terrestrial wildlife monitoring by enabling efficient, large-scale, long-term data collection with minimal disturbance (Sugai et al., 2019). Further, advances in low-cost hardware and automated processing have expanded its accessibility and scalability (Bradfer□Lawrence et al., 2024; Hill et al., 2018; Sugai et al., 2019). Seabirds are particularly well suited to PAM applications due to their high vocal activity (Nelson and Baird, 2001) and dense colonial nesting, producing substantial acoustic activity throughout the breeding season. Acoustic recorders can be deployed continuously, enabling long-term monitoring while mitigating accessibility challenges associated with remote offshore breeding colonies (Shonfield and Bayne, 2017; Van Doren et al., 2023). As a result, PAM is increasingly used in seabird research, including tracking within-colony flight patterns, estimating abundance, and documenting diel patterns in vocal behavior (Arneill et al., 2020; Colombelli□Négrel, 2023; Podolskiy et al., 2024).

While PAM has been successfully used to study phenology in passerines (Buxton et al., 2016; Van Doren et al., 2023), its application to seabird phenology remains limited. For passerines, a common PAM approach for tracking breeding phenology is quantifying vocal activity rate (VAR), or the number of calls produced within a given time period, which varies with breeding stage as birds arrive, court, and rear chicks (Jahn et al., 2017). In seabirds, VAR has primarily been used to assess colony occupancy, nest density, and abundance (Borker et al., 2014; Buxton and Jones, 2012; Zhao et al., 2022), despite evidence that acoustic activity may align better with breeding stages than nest density (Arneill et al., 2020). However, quantifying vocal activity in large, dense colonies is challenging due to overlapping vocalizations, even with modern machine-learning tools such as BirdNET (Kahl et al., 2021), and manual annotation would require substantial effort (Borker et al., 2014). As a result, VAR is difficult to apply at the colony scale, highlighting the need for PAM methods capable of efficiently quantifying vocal activity in colonies exceeding 100 nests or 0.63 nests/m^2^ (Borker et al., 2014).

When extracting individual calls becomes impractical, acoustic indices offer an automated alternative for processing acoustic datasets. Acoustic indices quantify temporal and spectral patterns in recordings, from simple summaries to complex estimates, enabling rapid processing of large datasets (Bradfer□Lawrence et al., 2023). Properly applied, acoustic indices can characterize soundscape patterns, serve as proxies for ecological activity and biodiversity (Bradfer□Lawrence et al., 2023), and estimate population size (Kloepper et al., 2016). However, the performance of acoustic indices across species and habitats remains poorly understood, particularly in seabird colonies where applications remain relatively limited. Nevertheless, several studies have shown promise for assessing colony dynamics, including nest density, in-colony flight activity, foraging behavior, and seasonal acoustic patterns (Arneill et al., 2020; Brownlie et al., 2020; Podolskiy et al., 2024), highlighting their potential as indicators of colony phenology.

Selecting an appropriate acoustic index requires aligning index design with the biological system to avoid misinterpretation (Bradfer□Lawrence et al., 2023). Indices developed to estimate species richness (e.g., Bioacoustic Index; Boelman et al., 2007) are poorly suited for single species aggregations, while others (e.g., ACI; <u>Pieretti et al., 2011</u>) can be difficult to interpret due to their computational complexity (Bradfer□Lawrence et al., 2023). In contrast, acoustic energy provides a simple, interpretable measure of amplitude over time, alleviating the need for detailed knowledge of spectral structure in the soundscape that is necessitated by more complex indices. While VAR may be conceptually ideal, it is often impractical due to high colony density and extensive call overlap. Prior studies in bats have demonstrated that acoustic energy can reflect the number of individuals in breeding colonies (Eddington et al., 2025; Kloepper et al., 2016), suggesting that colony-level amplitude may be a suitable proxy for VAR without needing to extract individual calls. Accordingly, acoustic energy represents a promising and scalable metric for tracking breeding phenology in colonial seabirds.

Tracking seabird phenology is especially important in polar regions where rapid ocean warming and sea-ice loss are altering breeding habitat and food availability (Cimino et al., 2023). Among Antarctic species, the Adélie penguin (*Pygoscelis adeliae)* is a well-established climate change sentinel (Ainley, 2002) with a tightly synchronized breeding period characterized by distinct behavioral stages, including egg laying, incubation, guard, post-guard, and fledge (Black, 2016; Spurr, 1975a, 1975b). Another colonial seabird species threatened by climate change is the common tern (*Sterna hirundo*). Common terns are migratory seabirds with a global distribution that disperse widely during winter foraging (Bugoni et al., 2005), but aggregate during the summer breeding season to form large, dense, and highly synchronized breeding colonies reaching thousands of pairs (Burger et al., 1988; Hernández-Matías et al., 2003). Like Adélie penguins, breeding is characterized by distinct phenological stages, including courtship, laying and incubation, hatching, and fledging associated with stage-specific behavioral shifts defined by parent-offspring interactions (Arnold et al., 2020).

In both Adélie penguins and common terns, breeding stages are not only defined by distinct behavioral states but also by predictable shifts in vocal behavior, positioning PAM as a promising tool for tracking breeding phenology in colonial seabirds. In Adélie penguins, adults engage in pair formation and copulation upon arrival at the colony (Black, 2016; Spurr, 1975a). During incubation, adults alternate nest duties while the other forages at sea (Spurr, 1975b), reducing colony attendance. Vocal activity during this stage is associated primarily with nest exchanges and aggressive interactions with conspecifics or predators (Black, 2016; Spurr, 1975a). During the guard stage, hatched chicks produce begging calls, while both adults and chicks produce mutual display calls associated with nest-duty exchanges and early vocal development (Spurr, 1975a, 1975b). Adults continue alternating nest duties during the guard stage until both eventually leave the chick to forage to meet rising energetic demands (Spurr, 1975b). As colonies transition to the post-guard stage, chick vocalizations become longer and more frequent (Adams et al., 2026), and unattended chicks aggregate into crèches (Black, 2016; Davis, 1982). Chicks continue to expand their vocal repertoire (Adams et al., 2026), while mobile chicks may also elicit aggressive territorial responses from nearby adults (Spurr, 1975b). As fledging approaches, parental provisioning declines in the two weeks preceding colony departure (Spurr, 1975b), and colony vocal activity is expected to decrease rapidly as chicks molt and fledge synchronously from colonies in early February (Cimino et al., 2023).

Common terns similarly exhibit a diverse vocal repertoire linked to breeding behavior that changes across the season. During courtship and pre-incubation, females produce begging calls indicating receptiveness to mate, while males produce copulation calls during pre-copulatory displays (Arnold et al., 2020). Terns also produce distinct vocalizations associated with territorial defense and aggressive interactions with conspecifics, behaviors that are especially frequent during nest establishment (Arnold et al., 2020; Cabot and Nisbet, 2013; Veen, 1987). Mating activity peaks surrounding egg laying and declines rapidly as birds transition to incubation (Blanchard and Morris, 1998; Wiggins and Morris, 1988). The incubation stage is associated with low-intensity brooding calls, with one adult attending the nest while the other forages at sea (Burger and Gochfeld, 1998; Nisbet and Cohen, 1975). Chick vocalizations begin shortly before hatching and develop rapidly through the first week post-hatch (Arnold et al., 2020). During the hatching stage, adults continue brooding and guarding chicks (Burger and Gochfeld, 1998), while territorial defense intensifies as chicks become increasingly mobile and adults respond aggressively to territorial intrusions (Quinn et al., 1994; Sorokaitė, 2005). Adults continue producing brooding calls until chicks fledge approximately one month after hatching (Burger and Gochfeld, 1998). During fledging, juveniles begin flying, gradually increasing the duration of off-colony flights as they begin foraging (Burger and Gochfeld, 1998), and develop flight calls (Arnold et al., 2020). As colonies transition to post-breeding dispersal, when birds depart the colony and prepare to migrate to overwintering grounds in early August (Caldwell et al., 2025), vocal activity is expected to diminish rapidly.

Despite differences in ecology and geography, both Adélie penguins and common terns share key features of colonial breeding. The timing of seabird breeding stages is strongly influenced by climate change (Carloni, 2018; Cimino et al., 2016; Dobson et al., 2017). In Adélie penguins, changes in sea-ice extent have altered breeding onset, productivity, and juvenile survival (Cimino et al., 2016; Dugger et al., 2014; Ninnes et al., 2011). Common terns exhibit both advances (Dobson et al., 2017; Ezard et al., 2007) and delays (Carloni, 2018) in lay dates in response to changing environmental conditions in the eastern and western hemispheres, respectively. These patterns underscore the importance of accurately characterizing phenology in colonial seabirds as environmental change continues to alter the timing and success of reproduction. Large colony sizes, synchronized breeding, and stage-specific vocal behaviors make both species well-suited for developing efficient, minimally invasive PAM methods to monitor phenology trends.

Here, we present a comparative case study examining the relationship between acoustic energy and breeding phenology in Adélie penguins and common terns. We hypothesize that colony-level acoustic energy will fluctuate predictably across phenological stages, with periods characterized by high social interaction and territoriality (e.g., courtship, chick-rearing, fledging) exhibiting increased amplitude relative to stages of reduced social interaction (e.g., incubation, post-breeding dispersal). To assess the efficacy of acoustic energy in studying seabird colonies, we (1) evaluated relationships between colony amplitude and phenological stage with generalized linear mixed-effects models, and (2) characterized seasonal patterns in colony soundscape across the breeding cycle while accounting for environmental covariates. By characterizing colony soundscapes using acoustic indices, this study explores the potential of PAM as a scalable tool for tracking breeding phenology in dense seabird colonies.

## 2. Methods

### 2.1. Study Site and Data Collection

#### 2.1.1. Adélie Penguin Colony on the Western Antarctic Peninsula

We collected data from Adélie penguin sub-colonies on Torgesen and Humble Islands near Palmer Station on the Western Antarctic Peninsula Figure S.1.). We deployed Wildlife Acoustics Song Meter Minis (SMM) across five sub-colonies from November 27, 2022, to February 27, 2023, during the Adélie penguin breeding season. Deployment occurred after egg laying to reduce nest abandonment risk during equipment setup (Giese, 1996). Recorders sampled five minutes per hour at a 24 kilohertz (kHz) sampling rate. We used data from four functioning rigs across four sub-colonies that recorded data without failure (Figure S.1.; Supplementary Methods S.1.1.1.). We obtained environmental data from the Palmer Automatic Weather Station (PAWS) at Palmer Station, Antarctica, approximately one kilometer from Humble and Torgersen Islands (Figure S.1.; Palmer Station Research Associate 2023).

We collected phenology data at the largest sub-colonies on each island and estimated colony-level phenology using mean egg-one lay and hatch dates (Supplementary Methods S.1.1.1.). For analyses, we defined incubation as the period between the mean lay date and the mean hatch date, guard as the period from the mean hatch date to 20 days after (Chapman et al., 2011; Cimino et al., 2023), and post-guard as the period from 21 days after the mean hatch date until chick fledging began We defined the fledge stage as the period when colonies started to disband to the end of the study (February 27, 2023), when chicks vacated colonies. Because breeding phenology is relatively synchronous within islands, we applied the same phenological stage boundaries to all three Torgersen sub-colonies.

#### 2.1.2. Common Tern Colony on Seavey Island, NH

We collected data from the common tern colony on Seavey Island, New Hampshire (Figure S.2.). Recorder locations were selected using historic census data to target areas primarily occupied by common terns (Figure S.2.; Supplementary Methods S1.1.2.) We deployed AudioMoth recorders (v. 1.0.0-1.2.0, Open Acoustic Devices, Southampton, UK) across 12 sites from May 11 to August 31, 2024. Recorders sampled 10 minutes per hour at a 48 kHz sampling rate on a continuous 24-hour schedule. Environmental covariates, including temperature, wind speed, precipitation, and moon illumination, were obtained from nearby monitoring stations (Supplementary Methods S.1.1.2).

During the summer of 2024, we monitored phenology and estimated colony-level phenology using mean egg-one lay and hatch dates (Supplementary Methods S.1.1.2.). We defined breeding stages using observed phenology and published life history estimates. The mean lay date of monitored nests was May 23, and the mean hatch date was June 17. Courtship spanned from recorder deployment to the day before the mean lay date (May 11 to 22), and incubation from the mean lay date to the day before the mean hatch date (May 23 to June 16). We defined the hatch stage as the period from the mean hatch date to 26 days post-hatch (June 17 to July 12), based on the median duration from hatch to fledge (Burger and Gochfeld, 1991; Nisbet and Drury, 1972). We defined the fledge stage as the period from the end of the hatch stage to the onset of dispersal (July 13 to 29). We defined the start of post-breeding dispersal (hereafter, the dispersal stage) as July 29, based on mean last detection dates from concurrent GPS tracking data (n=20; <u>Caldwell et al., 2025</u>), consistent with the reported average interval between fledging and family dispersal (15 days; Nisbet 1976). Manual inspection of acoustic data indicated that the last common tern vocalization detected occurred on August 17. Thus, we defined the dispersal stage as July 30 to August 16, and the post-dispersal period as August 17 to 31.

### 2.2. Acoustic Data Acquisition

We processed all acoustic data in R (R Core Team, 2025) using the seewave (Sueur et al., 2008) and tuneR (Borg, 2016) packages. For Adélie penguins, we excluded recordings degraded by high wind using an automated classifier and filtered recordings to the chick vocalizations range (Supplementary Methods S.1.2.1). For common terns, recordings were bandpass filtered to isolate species specific vocal frequencies and reduce geophonic noise (Supplementary Methods S.1.2.2). We calculated root-mean-square (RMS) power (relative [rel.] dB) in one-minute time windows using the rms() function in seewave (Sueur et al., 2008). RMS power quantifies acoustic energy as the root-mean-square of signal pressure over time (Madsen, 2005), with higher values indicating louder recordings. Because absolute sound pressure levels were not available for all devices (Supplementary Methods S.1.3.), we used relative RMS power calculated as:

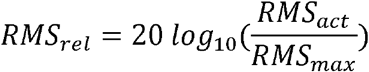

where RMSrel equals the RMS power measurement in relative decibel units, RMSact equals the recorded signal RMS power measurement, and RMSmax equals the RMS power of the maximum signal energy measurement possible (i.e., a one-second simulated floating-point file consisting of alternating −1 and +1 values).

### 2.3. Statistical Analyses

We conducted all statistical analyses in R (R Core Team, 2025). To assess differences in RMS power (rel. dB) among breeding stages, we fit a general linearized mixed-effects model (GLMM) with a Gamma error distribution and log link. We rescaled negative RMS power values relative to the colony-specific sound floor and added a small constant to transform negative values and avoid zeros (Supplementary Methods S.1.4.). We included RMSscaled as the response and phenological stage as a categorical fixed effect. To account for diel structure in acoustic activity, we included sine and cosine transformations of the hour of the day. We excluded day-of-year terms because they absorbed variation associated with phenological stage. Environmental covariates known to effect both seabird behavior and acoustic signal propagation included wind speed and temperature in both systems, with precipitation and moon illumination included only in the common tern model (<u>Kasten et al., 2012; Pérez□Granados and Schuchmann, 2021</u>; Supplementary Methods S.1.4.). We included recorder as a random intercept to account for repeated measurements and variation in device sensitivity.

We estimated marginal means and conducted post-hoc comparisons between consecutive phenological stages (Supplementary Methods S.1.4.). We converted model coefficients and post-hoc estimates to percent change to facilitate interpretation in the text, while original values were reported in the tables. All effects represented model-estimated changes in mean RMS power with other covariates held constant. We treated courtship as the reference stage for common terns and incubation for the Adélie penguins; all percent changes were interpreted relative to these reference stages for the GLMM interpretations. Post-hoc comparisons quantified differences between consecutive stages and were interpreted independently of the reference stage.

## 3. Results

### 3.1. Phenological Effects

For both species, scaled RMS power varied significantly across phenological periods, diel cycles, and environmental conditions (Figure 1; Tables S.1. & S.2.). For the Adélie penguins, phenological stage was a strong predictor of variation in acoustic energy (Figure 1A-B; Table S.1.). Holding diel and environmental covariates constant, the model estimated that mean RMS power increased during the guard (+12.4%) and post-guard (+27.6%) stages relative to incubation, before declining during the fledge stage (-32.1%) as penguins dispersed (*p*<0.0001; Figure 1A-B; Table S.1.). Post-hoc comparisons between consecutive stages further clarified these transitions, indicating that model-estimated mean RMS power increased from incubation to guard (+12.4%) and from guard to post-guard (+13.5%), followed by a sharp decline from post-guard to fledge (-46.8%; *p*<0.0001; Table S.3.).

**Figure 1.**
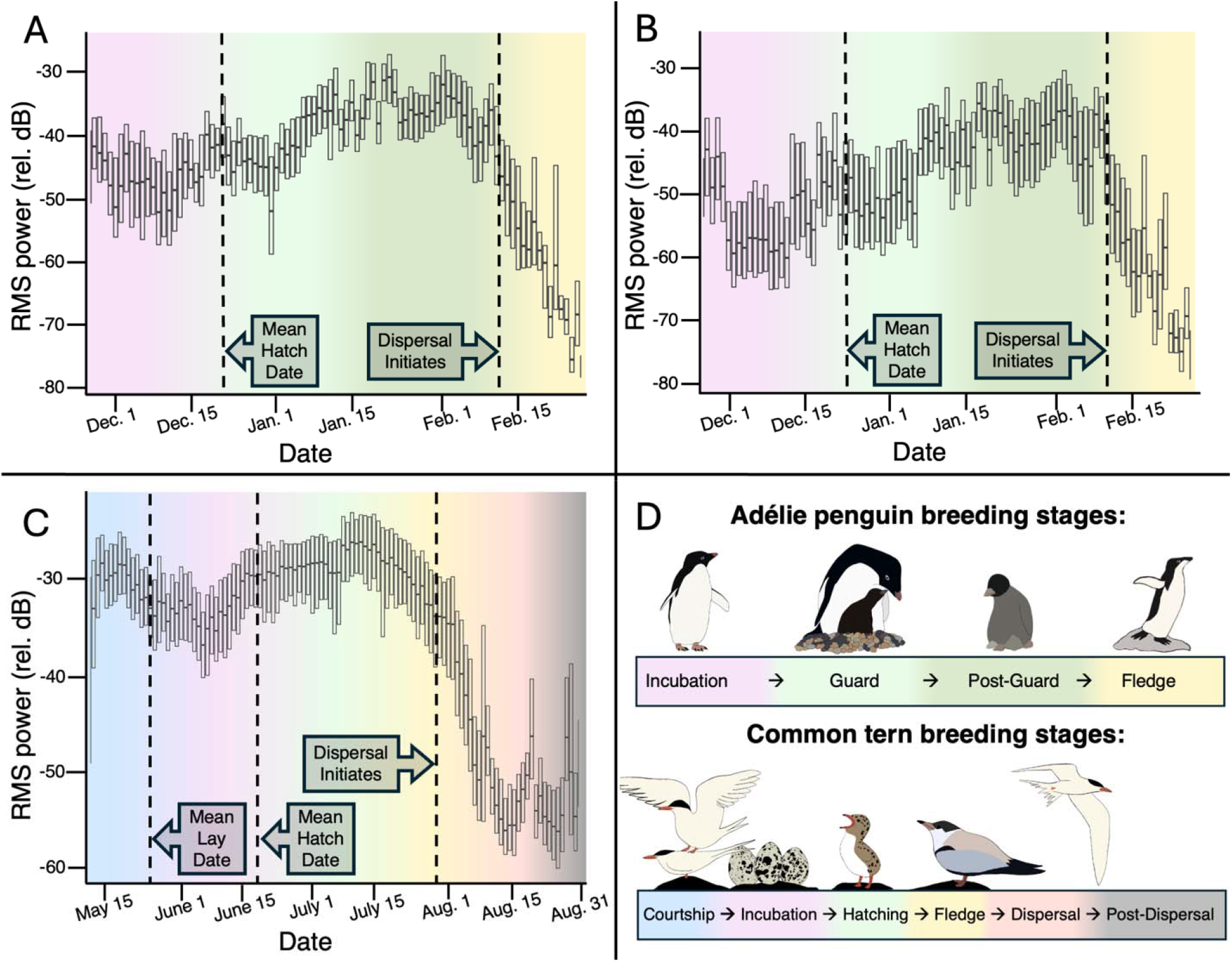
Seasonal variation in RMS power (relative dB) across the breeding season in two Adélie penguin colonies and one common tern colony. RMS power is shown from late-November 2022 to late-February 2023 in the Adélie penguin colony on (A) Humble Island, Antarctica, and (B) Torgersen Island, Antarctica, and (C) RMS power is shown from mid-May to late-August 2024 in the common tern colony on Seavey Island, NH, USA. Boxplots show interquartile ranges with median values indicated by solid black lines in the center of the boxes. Vertical dashed lines denote key phenological observations within the recording period, and shading indicates breeding stages. (D) The breeding stage legend corresponds to the shaded regions in panels A-C.

Similarly, phenological stage strongly predicted changes in estimated mean acoustic energy in the common tern colony (Figure 1C; Table S.2.). Relative to courtship and holding all other predictors constant, estimated mean RMS power decreased significantly during incubation (-12.2%), hatching (-9.93%), fledge (-4.55%), with substantial decreases during dispersal (-47.3%) and post-dispersal (-57.1%) as birds left the colony (*p*<0.0001; Figure 1C; Table S.2.). Post-hoc comparisons between consecutive stages clarified these transitions, indicating a decline in model-estimated mean RMS power from courtship to incubation (-12.2%), followed by increases from incubation to hatching (+2.5%), and from hatching to fledge (+6%), before a sharp decline during dispersal (-44.8%) and a continued decrease into the post-dispersal stage (-18.6%; *p*<0.0001; Table S.4.).

### 3.2. Environmental Covariates, Diel Patterns, and Random Effects

Environmental conditions significantly influenced the estimated mean RMS power in both systems (Tables S.1. & S.2.). Increased wind speed was associated with small but significant increases in estimated mean RMS power in both systems (<2%; *p*<0.001; Tables S.1. & S.2.). Higher temperatures were also positively associated with RMS power, increasing mean RMS power by 0.9% and 3.57% in the tern and penguin models, respectively (*p*<0.001; Tables S.1. & S.2.). In the common tern model, rainfall increased estimated mean RMS power by 3.25% compared to dry conditions, while increased moon illumination was associated with a 13.8% decrease in estimated mean RMS power compared to darker conditions (*p*<0.0001; Table S.2.). Diel variation indicated pronounced within-day patterns in acoustic activity in both models (*p*<0.001; Tables S.1. & S.2.). Between-recorder variability was low across datasets (common tern: random intercept SD = 0.014; Adélie penguin = 0.033), indicating phenological, temporal, and environmental predictors explained most variation in estimated mean RMS power across sites.

## 4. Discussion

Our findings support that colony-wide acoustic energy closely tracks breeding phenology in two species of colonial seabird, the Adélie penguin and the common tern. Across both systems, distinct shifts in acoustic energy correspond to major transitions in colony behavior, including courtship, incubation, chick rearing, fledging, and post-breeding dispersal. These patterns suggest that PAM can capture biologically meaningful changes in colony activity without requiring direct observation of phenological indicators (e.g., egg laying, hatching) that introduce potentially harmful investigator disturbance to breeding seabirds (Carey, 2009; Carney and Sydeman, 1999). Despite substantial differences in colony structure, geography, and breeding ecology between the two species, similar stage-specific changes in acoustic energy emerged, indicating that soundscape dynamics may provide a broadly applicable indicator of breeding phenology in colonial seabirds.

### 4.1. Adélie Penguins

At the Adélie penguin colonies, recorders were deployed after egg laying to minimize nest-abandonment associated with investigator disturbance. Acoustic energy was lowest during incubation, likely reflecting reduced colony attendance associated with alternating nest duties (Black, 2016; Spurr, 1975b). As chicks began hatching, acoustic energy increased during the guard stage, aligning with the onset of parent-offspring interactions and chick vocal development (Spurr, 1975a, 1975b). Acoustic energy continued to rise and peaked during the post-guard stage, coinciding with intensified parental provisioning, chick mobility, and crèche formation (Adams et al., 2026; Black, 2016; Davis, 1982; Spurr, 1975b, 1975a). Elevated acoustic energy during this stage likely reflected increased chick vocalizations and conspecific interactions throughout the colony (Adams et al., 2026; Spurr, 1975b). Acoustic energy then declined sharply during the fledge stage as parent-offspring interactions decreased and colonies dispersed in early February (Spurr, 1975b). The stage-specific shifts in colony activity aligned with predictable changes in RMS power (Figure 1A-B), supporting the use of colony soundscape as an indicator of breeding phenology in Adélie penguins.

### 4.2. Common Terns

For common terns, acoustic energy was high in mid-May during pair formation, courtship, and nest establishment as birds arrived at the colony (Burger and Gochfeld, 1998). This period is characterized by frequent territorial interactions and mating behavior (Arnold et al., 2020; Cabot and Nisbet, 2013), which likely contributed to elevated acoustic energy. Consistent with the decline in mating activity following egg laying (Blanchard and Morris, 1998; Wiggins and Morris, 1988), RMS power decreased as birds transitioned to a three-week incubation period associated primarily with low-intensity brooding calls (Burger and Gochfeld, 1998; Nisbet and Cohen, 1975; Zogby et al., 2026). The peak and subsequent decline in acoustic energy as birds transitioned from courtship and laying to incubation represented a defining transition in the breeding season soundscape of common terns, aligning with known phenological patterns in colony activity associated with courtship and mating.

Following this decline, acoustic energy increased later in the incubation stage in the week preceding the mean hatch date and remained elevated during the hatching stage, coinciding with the onset of parent-offspring interactions, chick vocal development, and heightened territorial defense within the colony (Arnold et al., 2020; Burger and Gochfeld, 1998; Quinn et al., 1994; Sorokaitė, 2005). Acoustic energy continued to increase into the early fledge stage, consistent with the period when juveniles developed flight calls and began flying at the colony (Arnold et al., 2020; Burger and Gochfeld, 1991; Nisbet and Drury, 1972). Later in the fledge stage, as fledglings progressively increased flight duration and initiated foraging, time spent off colony was expected to increase (Arnold et al., 2020), corresponding with the observed decline in RMS power. Finally, as birds departed the colony during post-breeding dispersal in late July, acoustic energy declined rapidly, reaching seasonal lows by mid-August. These stage-specific patterns in colony activity corresponded with predictable shifts in acoustic energy (Figure 1C), demonstrating that colony soundscape can serve as a reliable proxy for breeding phenology in common terns.

Across both species, acoustic energy followed a consistent seasonal trajectory characterized by elevated RMS power during periods of high social and reproductive activity and reduced RMS power during incubation. This was followed by pronounced increases during chick-rearing stages and then declines associated with fledging and colony dispersal. While the specific behavioral drivers differed somewhat between systems, these parallel patterns indicated that colony soundscape dynamics reflected shared temporal patterns in breeding phenology in colonial seabirds. These predictable shifts in acoustic energy in both common terns and Adélie penguins suggested that PAM could provide a scalable method for identifying key breeding transitions in colonial seabirds.

### 4.3. Environmental Effects

In terms of environmental effects, moonlight exhibited a negative relationship with acoustic energy in the common tern colony, consistent with prior work showing reduced nocturnal vocal activity in seabirds during increased lunar illumination (Linares et al., 2022; Mougeot and Bretagnolle, 2000). Manual inspection of spectrograms from rainy recordings revealed broadband interference from raindrops hitting the recorder within the 1-12 kHz frequency range used in our analyses, suggesting that the positive relationship between rainfall and acoustic energy reflects abiotic noise rather than increased biological activity. In both the common tern and Adélie penguin datasets, higher wind speed was associated with small increases in acoustic energy. Although filtering reduced low-frequency geophonic noise in both datasets, some wind noise during high wind periods likely exceeded the filtering threshold (Kasten et al., 2012), particularly under the extreme wind conditions experienced at the Adélie penguin colonies (Fradet et al., 2025).

Controlling for stage-dependent effects, higher temperatures were also associated with increased acoustic energy across both systems. Although the effects of temperature on avian vocal behavior vary across species and contexts (Cordonnier et al., 2023; Pérez□Granados and Schuchmann, 2021), higher temperatures are generally associated with reduced calling behavior due to heat stress (Soravia et al., 2021). In our study, however, temperatures may not have exceeded behavioral thresholds that suppress vocal activity (common tern colony: mean = 19.3 °C, S.D = 3.9, max = 31.7), particularly for the Adélie penguin colonies where temperatures remained low with generally limited variability (mean = 2 °C, S.D. = 1.7, max = 8.6). Moreover, the effect size of temperature in the common tern dataset was minimal (0.13%), suggesting a weak or negligible influence on vocal behavior. For Adélie penguins, warmer temperatures may correspond to more favorable weather conditions within an otherwise harsh environment, potentially facilitating increased conspecific interaction and communication. Together, these findings highlight the importance of accounting for environmental influences when interpreting biological soundscapes, given the close links between environmental conditions, seabird behavior, and the timing of breeding phenology.

### 4.4. Implications, Future Research, and Conclusion

To our knowledge, this study is the first to evaluate acoustic energy as a tool for monitoring seabird breeding phenology. Previous studies have examined other acoustic indices such as the Acoustic Complexity Index and the Bioacoustic Index for monitoring seabird breeding colonies (Arneill et al., 2020; Brownlie et al., 2020), but these metrics rely on complex quantifications of the time- and frequency-domains that can be difficult for non-acousticians to interpret. Furthermore, VAR is conceptually well-suited for tracking breeding phenology, but extracting individual calls from dense seabird colonies remains impractical at large spatial scales (Arneill et al., 2020; Borker et al., 2014; Brownlie et al., 2020). Building on prior work linking acoustic energy to population size in colonial bats (Eddington et al., 2025; Kloepper et al., 2016), our findings suggest that acoustic energy may provide a practical and intuitive measure of colony-level vocal activity in seabirds by linking phenological shifts in colony behavior directly to changes in RMS power.

Traditional seabird monitoring programs have revealed important drivers of breeding phenology and reproductive success (Cimino et al., 2023, 2019) but are often require limited in spatial scope, require extensive mounts of time, and can disturb breeding colonies (Carey, 2009; Carney and Sydeman, 1999; Cimino et al., 2016; Hinke et al., 2018; Shonfield and Bayne, 2017). Such challenges are especially important in rapidly changing systems, including the Western Antarctic Peninsula and the Gulf of Maine (Cimino et al., 2016; Hinke et al., 2018; Pershing et al., 2021). Our results suggest that PAM using acoustic energy may provide a scalable, minimally invasive approach for tracking breeding phenology in seabird colonies, complementing traditional monitoring programs by extending phenological observations across more colonies and longer time periods.

Future research could investigate how acoustic energy relates to colony size, attendance, and reproductive success, and evaluate the extent to which these relationships hold across additional years, species, and ecosystems. Establishing explicit relationships between acoustic energy and VAR could further strengthen the applicability of this approach, as VAR has been more widely explored in seabird monitoring. Additional work examining how environmental variables influence seabird vocal behavior in breeding colonies would also improve the interpretation of PAM datasets and may provide insight into how climate change affects these behaviors. Together, these efforts would help refine the application of soundscape metrics for long-term seabird monitoring.

Overall, our results demonstrate that acoustic energy provides a simple yet informative measure of colony-level activity that reflects predictable behavioral changes across the breeding season. In both common tern and Adélie penguin colonies, transitions between breeding stages produced consistent shifts in soundscape amplitude driven by changes in mating behavior, territorial interactions, chick development, and parent-offspring communication. These findings indicate that colony soundscapes can capture key phenological transitions in large seabird colonies, highlighting the potential of PAM as a practical tool for tracking breeding dynamics in remote or rapidly changing environments.

## CRediT authorship contribution statement

**Valerie M. Eddington:** Conceptualization, methodology, software, data curation, formal analysis, funding acquisition, investigation, writing – original draft, writing – review & editing, visualization. **Danielle T. Fradet:** Conceptualization, methodology, software, data curation, formal analysis, investigation, writing – original draft, writing – review & editing. **Elizabeth C. Craig:** Conceptualization, methodology, resources, investigation, writing – review & editing, supervision, project administration, funding acquisition. **Megan A. Cimino:** Conceptualization, methodology, resources, investigation, writing – review & editing, supervision, project administration, funding acquisition. **Easton R. White:** Conceptualization, methodology, writing – review & editing, supervision, funding acquisition. **Laura N. Kloepper:** Conceptualization, methodology, writing – review & editing, supervision, funding acquisition

## Funding

This research was funded by the UNH School of Marine Science and Ocean Engineering Chase Family Peer Mentorship Fund, the UNH Graduate School Summer Teaching Assistant Fellowship, and National Science Foundation EAGER Grant No. 2026045 and Award No. 2226886. The Isles of Shoals Tern Conservation Program is supported by the New Hampshire Fish and Game Department’s Nongame and Endangered Wildlife Program, the United States Fish and Wildlife Service (New Hampshire State Wildlife Grants), and the Shoals Marine Laboratory.

## Declaration of Competing Interests

The authors have no relevant financial or non-financial interests to disclose.

## Acknowledgements

We would like to thank Aliya Caldwell, Willow Dalehite, Orena Wong, Joe Brosseau, and Hope Caliendo for assistance in deploying and recovering AudioMoths and collecting field data. Our sincere thanks to Darren Roberts and Megan Roberts for the Adélie penguin data collection. Finally, we would also like to thank all of the previous and current members of the QMEL and EAB labs for their support, feedback, and encouragement.

Tern colony fieldwork was permitted under protocol #220304 approved by the University of New Hampshire’s Institutional Animal Care and Use Committee along with permissions from the NH Fish and Game Department and the US Fish and Wildlife Service for annual colony-monitoring activities. Adelie penguin work was approved by the Institutional Animal Care and Use Committee at UC Santa Cruz (protocol Cimim2204dn). This publication is contribution #219 to the Shoals Marine Laboratory.

## Data Availability

The data that support the findings of this study are available from the corresponding author upon reasonable request.

## Notes

### Competing Interest Statement

The authors have declared no competing interest.

