## Supplemental Materials for "Soundscape reflects breeding phenology in colonial seabirds"

**S.1. Methods**

**S1.1. *Study systems and data collection expanded***

**S1.1.1. *Adelie penguin colony on the Western Antarctic Peninsula***

We selected sub-colonies on Torgersen and Humble Islands to capture a range of sub-colony sizes (min: 15 nests; max: 517 nests). Dominant sounds in the soundscape included adult Adélie penguin vocalizations, chick vocalizations, and wind (Figure S.3). Recorders were configured with a six dB gain in a 16-bit depth; shorter recording windows were used to preserve battery life in extreme cold. We mounted SMMs at a height of one meter on PVC rigs and placed them at sub-colony edges with microphones facing inward. We deployed one rig per sub-colony except at the largest (>500 nests), where we deployed three. Elephant seals (*Mirounga angustirostris*) crushed two rigs, and high winds ripped the recorder from another. Thus, we excluded these three rigs from analysis due to lack of data.

At each sub-colony, we monitored six transect lines of five nests per line and recorded nest status daily (Cimino et al., 2019), including egg laying and chick hatching dates for each nest. We estimated colony-level phenology by averaging egg-one lay dates to obtain the mean lay date and egg-one hatch dates to obtain the mean hatch date. When nests checks were missed due to weather, we corrected estimated dates using the known incubation duration following Cimino et al. (2019). We excluded dates that could not be corrected (n=8)

**S.1.1.2 Common tern colony on Seavey Island, NH, USA**

The Seavey Island common tern colony hosts a long-term monitoring program. In the five years preceding this study, the mixed-species colony consisted of approximately 3,000 breeding pairs of common terns, ~100 breeding pairs of roseate terns (*S. dougallii*), and one breeding pair of Arctic terns (*S. paradisaea*)(Craig, 2024). Roseate terns nest near conspecifics, demonstrating a clustered distribution, whereas common terns are more evenly distributed across the island.

We programmed AudioMoths with low gain sensitivity. Two AudioMoth devices per site to minimize data gaps during servicing. Early-season data loss due to alkaline battery failure was resolved by switching to lithium batteries on June 16, 2024. Devices recorded 106 days on average. We mounted recorders ~1.25 meters above ground on custom PVC rigs and stabilized them with adhesive foam to reduce vibration. We obtained air temperature (°C) and wind speed (m/s) data from the NOAA weather station located on White Island, New Hampshire (Station IOSN3), and daily precipitation values from the Shoals Marine Laboratory located on neighboring Appledore Island, Maine (https://sustainablesml.org/pages/export.php). On days with precipitation, we manually verified rain presence in spectrograms. We obtained moon illumination data from the NOAA National Weather Service Weather Forecast Office in Norton, Massachusetts (https://www.weather.gov/box/sunmoon).

We monitored phenology at eight fenced productivity plots on Seavey Island spanning representative habitat variation across the island, encompassing variation in vegetative cover and plant community, topography, and exposure (mean area = 13.5 ± 2.5 m^2^, total monitored area = 108 m^2^). We recorded nest status every 1-2 days during egg laying and hatching and every 3-4 days thereafter, documenting egg-laying and chick-hatching dates for each nest. We estimated colony-level phenology by averaging egg-one lay and hatch dates of all monitored nests to obtain mean lay and hatch dates. Although some breeding pairs may renest following initial nest failure, initiating a secondary breeding schedule, for our analyses, we focused only on the primary breeding schedule of the colony (Arnold et al., 2020). In 2024, 96% of monitored nests followed the primary breeding schedule.

**S.1.2. *Acoustic Preprocessing***

**S.1.2.1. *Adelie Penguin Data***

Zhao et al. (2022) reported a significant relationship between wind speed and Adélie penguin chick vocalization rate and recommended removing wind-degraded recordings. The 2022-2023 breeding season was one of the windiest on record at Palmer Station, with PAWS data indicating peak gusts up to 60.6 knots and frequent gusts >40 knots, resulting in clipping and acoustic masking. To address irreparably degraded files, we developed a custom high-wind detector in BirdNET (Kahl et al., 2021) and applied it to our dataset (Fradet et al., 2025). We removed all five-minute recordings with a high-wind detection confidence score ≥ 0.56, excluding 857 recordings (9.6%) from analyses. We then applied a 2.5 to 5 kHz Butterworth bandpass filter using bwfilter() in the seewave package in R (Sueur et al., 2008), corresponding to the reported vocalization range of Adélie penguin chicks (Zhao et al., 2022).

**S.1.2.2. *Common Tern Data***

To isolate the primary frequency range of tern vocalizations (Zogby et al., 2026), reduce environmental noise below 1 kHz (Kasten et al., 2012), and minimize bias from inter-device variability above 12 kHz (Figure S.4), we applied a first-order Butterworth bandpass filter from 1 to 12 kHz using the seewave package in R (Sueur et al., 2008).

**S.1.3. *RMS (relative dB) calculation details***

We imported audio files in R as 16-bit linear pulse-code modulation (PCM) format (−32,768 and 32,768) and standardized them to double floating-point values (−1 to 1) to align with common bioacoustics analysis software (i.e., MATLAB [MathWorks, Natick, MA] and Raven Pro [Cornell Lab of Ornithology, Ithaca, NY]). We calculated root-mean-square (RMS) power (relative [rel.] dB) in one minute time windows, consistent with temporal aggregation used in previous seabird bioacoustic studies (Arneill et al., 2020; Borker et al., 2014; Brownlie et al., 2020; Oppel et al., 2014; Towsey et al., 2014). RMS power quantifies acoustic energy as the root-mean-square of signal pressure over time (Madsen, 2005), with higher values indicating louder recordings. Since AudioMoths do not come with reported sensitivity values from the manufacturer, we could not calculate amplitude values in sound pressure level, which is typically used to quantify amplitude (Lynch et al., 2011; Shannon et al., 2016). Instead, we calculated the relative RMS power of the signal values. To calculate RMS power in relative decibels for each minute of recording, we converted RMS signal power values to RMS relative power in relative decibels.

**S.1.4. Data Transformation and Statistical Analyses (expanded)**

We fit the model using the glmer() function in lme4 (Bates et al., 2015). The Gamma distribution with a log link was selected due to the continuous, right-skewed nature of RMS power data (Zuur et al., 2009). Because gamma models require non-zero positive data (Zuur et al., 2009), we rescaled negative RMS power values relative to the colony-specific sound floor and added a small constant to avoid zeros:

$${RMS}_{scaled}=\frac{{RMS}_{rel}-\min\left( {RMS}_{rel} \right)}{\max\left( {RMS}_{rel} \right)-\min\left( {RMS}_{rel} \right)}+0.000000000001$$

where RMSscaled is the RMS power (rel. dB) values rescaled relative to the colony-specific noise floor, and RMSrel is the measured RMS power in relative decibel units.

We initially considered including cyclic day-of-year terms; however, these terms absorbed variation associated with phenological stage and were excluded from final models. We excluded moon illumination from the Adélie penguin model because the Western Antarctic Peninsula experiences no true night during austral summer, and precipitation was not included due to limitations in data availability and relevant for the Antarctic system. We used the emmeans package (Lenth and Piaskowski, 2017) to estimate marginal means and conduct post-hoc comparisons between consecutive phenological stages. We evaluated model assumptions by inspecting residuals for distributional fit, homogeneity, and independence, and we used Akaike’s Information Criterion for model selection.

**Figure and Tables**
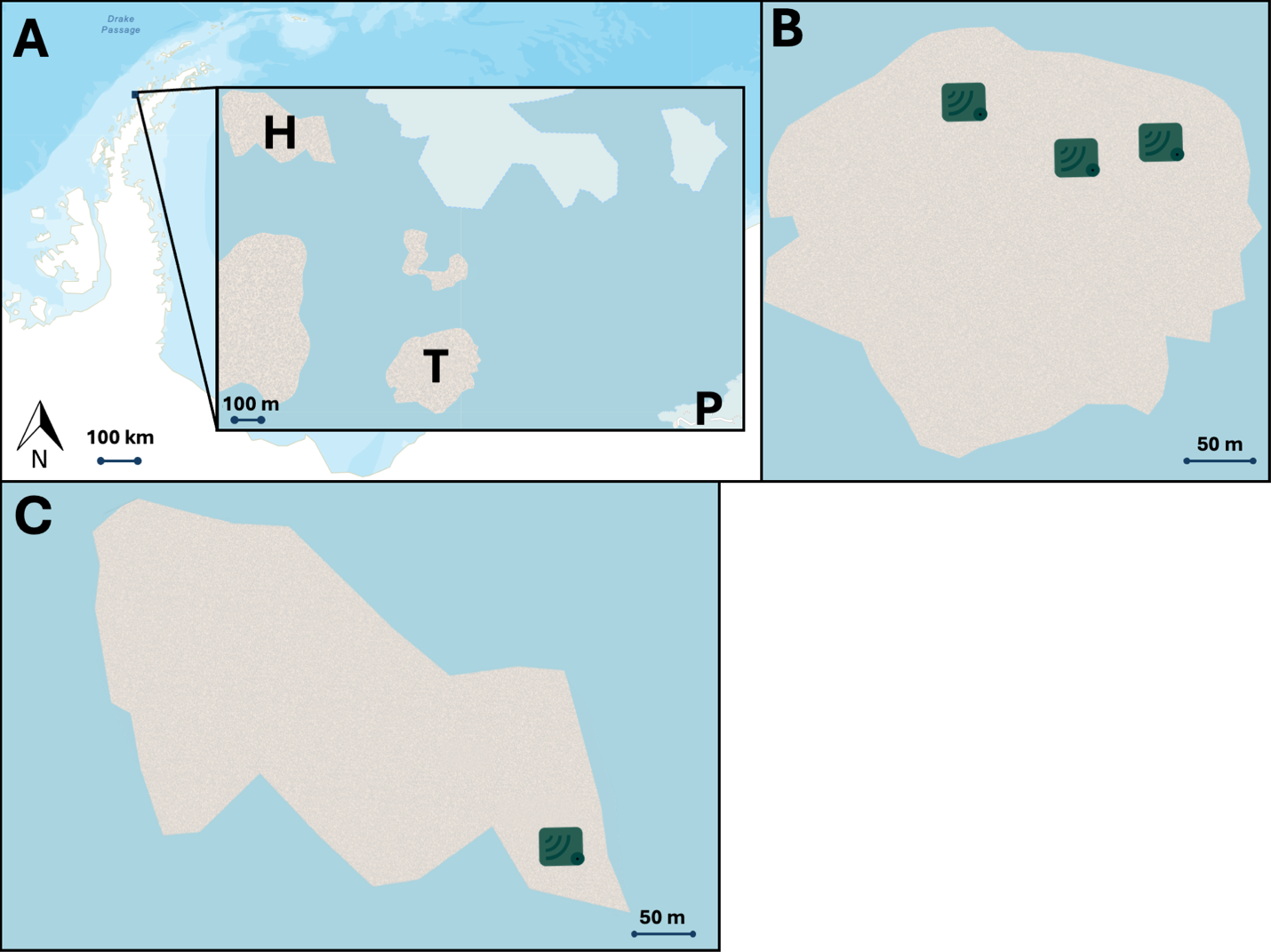
Figure S.1. A) The location of the Adélie penguin study sites within the Western Antarctic Peninsula. The inset maps show the relative location of Humble Island (H), Torgersen Island (T), and Palmer Station (P). B) Inset map of Torgersen Island. C) Inset map of Humble Island. For both B and C, green icons represent the location of audio recorders.

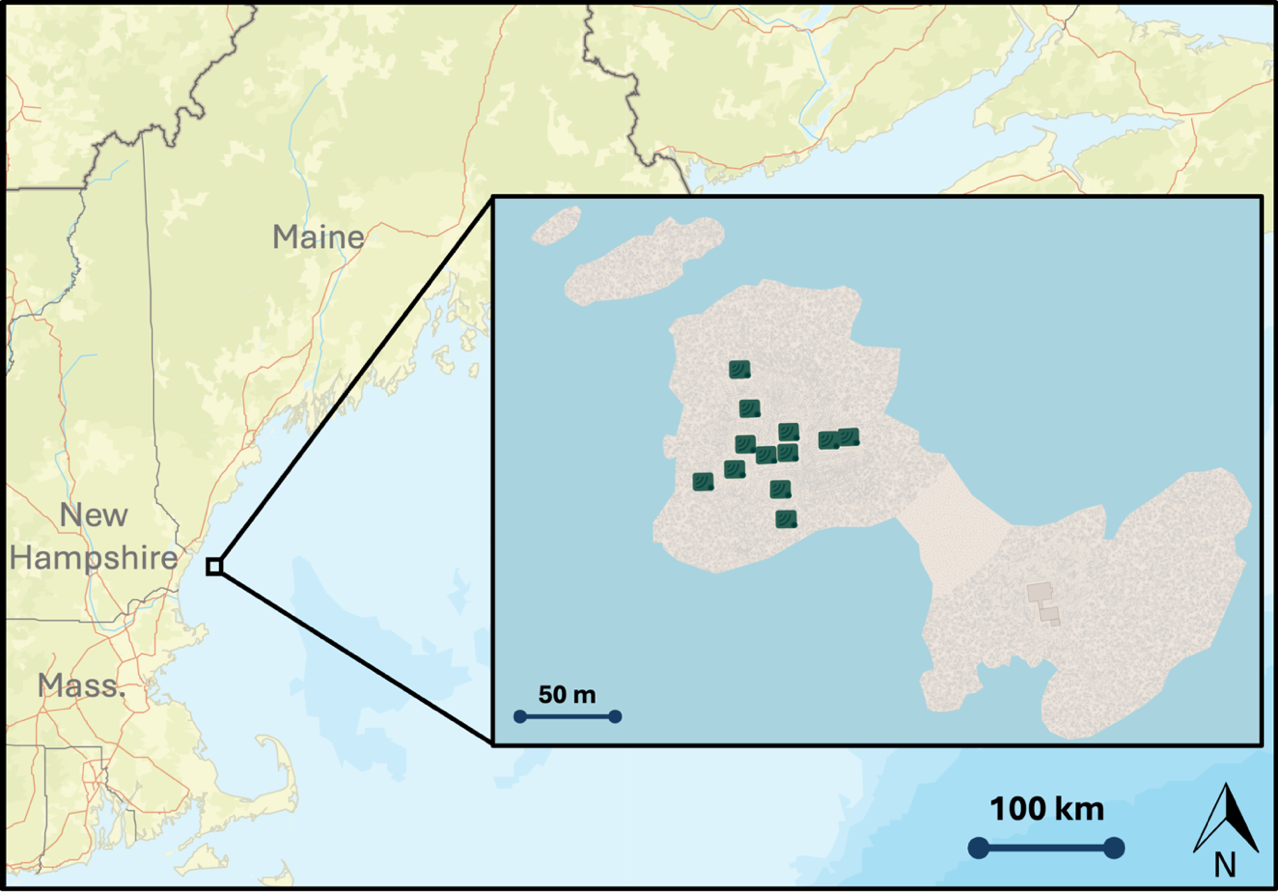

Figure 2. The inset map shows Seavey Island (left) and White Island (right), New Hampshire, located within the Gulf of Maine approximately six miles off the coast of New Hampshire. White and Seavey Islands are connected by a land bridge at low tide. Green icons on Seavey Island represent AudioMoths locations.

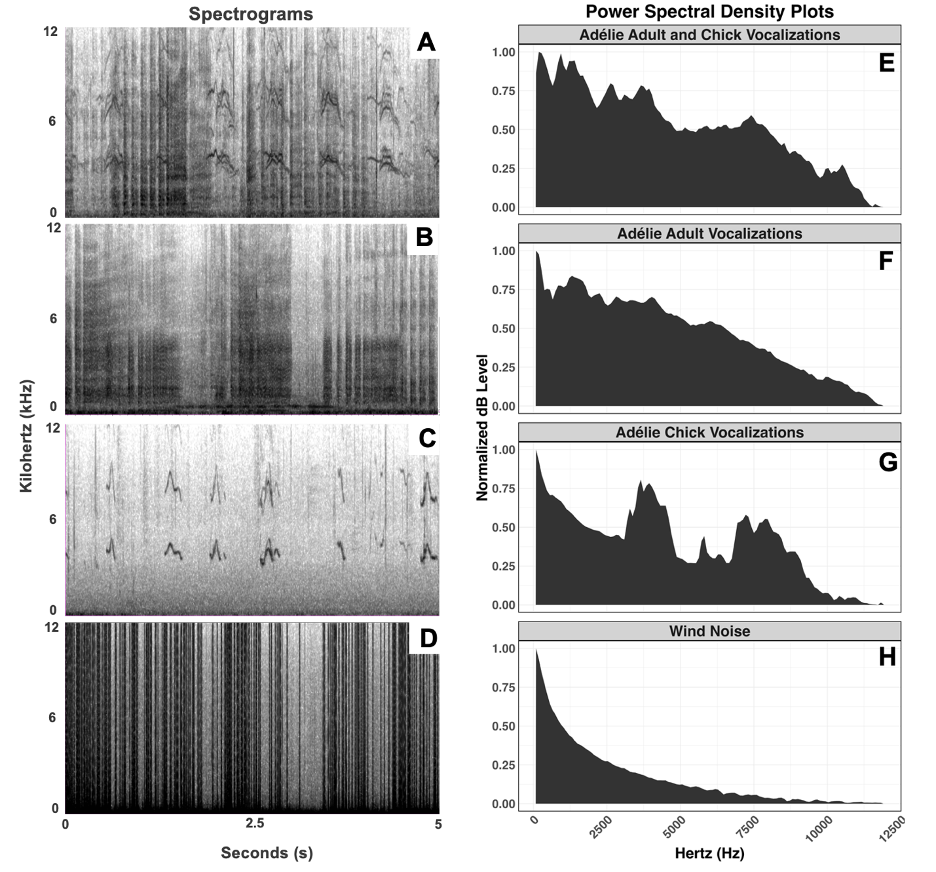

Figure S.1. Spectrograms of 5-second audio clips of A) Adélie penguin adult and chick vocalizations, B) adult vocalizations, C) chick vocalizations, and D) high wind noise. In spectrograms, darker colors represent higher amplitude sounds. Panels E-H are the corresponding normalized power spectral density plots of the same audio clips.

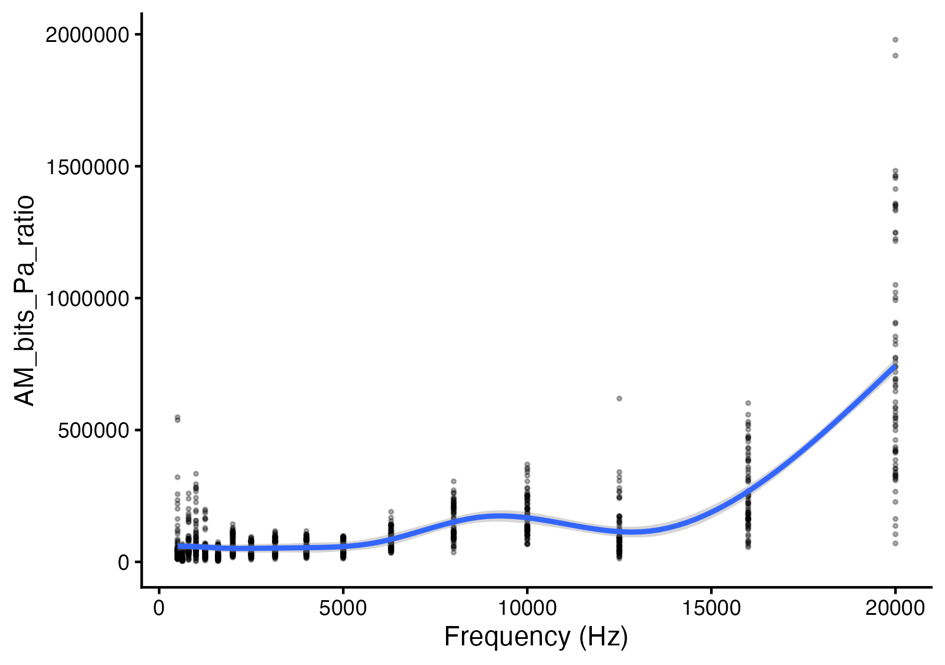

Figure S.4. The frequency response of AudioMoth devices from 500 Hz to 20,000 Hz.

Table S.1. Results of the GLMM analyzing the effect of phenological stage, temporal factors, and environmental covariates on RMS power at the Adélie penguin colonies in 2022 and 2023.

| **Effect** | **Estimate** | **Standard Error** | ***t*-value** | ***p*-value** |
| --- | --- | --- | --- | --- |
| Incubation (intercept) | -0.884 | 0.051 | -17.4 | <0.00001 |
| Guard | 0.117 | 0.0054 | 21.7 | <0.00001 |
| Post-Guard | 0.244 | 0.0047 | 51.8 | <0.00001 |
| Fledge | -0.387 | 0.0053 | -73.4 | <0.00001 |
| Cosine hour of day | -0.0370 | 0.0025 | -14.9 | <0.00001 |
| Sine hour of day | -0.00727 | 0.0026 | -2.79 | 0.00524 |
| Wind | 0.0125 | 0.00042 | 29.7 | <0.00001 |
| Temperature | 0.0351 | 0.0013 | 27.9 | <0.00001 |

Table S.2. Results of the GLMM analyzing the effect of phenological stage, temporal factors, and environmental covariates on RMS power at the common tern colony in 2024.

| **Effect** | **Estimate** | **Standard Error** | ***t*-value** | ***p*-value** |
| --- | --- | --- | --- | --- |
| Courtship (intercept) | 3.46 | 0.015 | 236 | <0.00001 |
| Incubation | -0.130 | 0.0025 | -49.3 | <0.00001 |
| Hatching | -0.105 | 0.0027 | -33.3 | <0.00001 |
| Fledge | -0.0465 | 0.0030 | -6.12 | <0.00001 |
| Dispersal | -0.640 | 0.0029 | -214 | <0.00001 |
| Post-Dispersal | -0.846 | 0.0026 | -315 | <0.00001 |
| Cosine hour of day | -0.0676 | 0.00083 | -83 | <0.00001 |
| Sine hour of day | 0.0211 | 0.00081 | 22.9 | <0.00001 |
| Wind | 0.00269 | 0.00002 | 136 | <0.00001 |
| Temperature | 0.00898 | 0.00021 | 33.8 | <0.00001 |
| Illumination | -0.148 | 0.0017 | 93.8 | <0.00001 |
| Rain | 0.0320 | 0.0045 | 5.08 | <0.00001 |

Table S.3. Results of the post-hoc comparisons of RMS power (rel. dB) between consecutive phenological stages for the Adélie penguin colony derived from estimated marginal means. Pairwise differences represent changes in estimated mean RMS power between adjacent stages while holding all other model covariates constant.

| **Contrast** | **Percent Change** | **Estimate** | **Standard Error** | **z** | ***p*-value** |
| --- | --- | --- | --- | --- | --- |
| Incubation 🡪 Guard | +12.4 | 0.117 | 0.0054 | 21.7 | <0.00001 |
| Guard 🡪 Post-Guard | +13.5 | 0.127 | 0.005 | 25.3 | <0.00001 |
| Post-Guard 🡪 Fledge | -46.8 | -0.631 | 0.0051 | -123 | <0.00001 |

Table S.4. Results of the post-hoc comparisons of RMS power (rel. dB) between consecutive phenological stages for the common tern colony derived from estimated marginal means. Pairwise differences represent changes in estimated mean RMS power between adjacent stages while holding all other model covariates constant.

| **Contrast** | **Percent Change** | **Estimate** | **Standard Error** | **z** | ***p*-value** |
| --- | --- | --- | --- | --- | --- |
| Courtship 🡪 Incubation | -12.2 | -0.127 | 0.0025 | -52.2 | <0.00001 |
| Incubation 🡪 Hatching | +2.53 | 0.0250 | 0.002 | 12.5 | <0.00001 |
| Hatching 🡪 Fledge | +5.98 | 0.0581 | 0.0019 | 30.5 | <0.00001 |
| Fledge 🡪 Dispersal | -44.8 | -0.594 | 0.0022 | -273 | <0.00001 |
| Dispersal 🡪 Post-Dispersal | -18.6 | -0.206 | 0.002 | -102 | <0.00001 |
